# Sustained Released of Bioactive Mesenchymal Stromal Cell-Derived Extracellular Vesicles from 3D-Printed Gelatin Methacrylate Hydrogels

**DOI:** 10.1101/2021.09.28.462252

**Authors:** Louis J. Born, Shannon T. McLoughlin, Dipankar Dutta, Bhushan Mahadik, Xiaofeng Jia, John P. Fisher, Steven M. Jay

**Affiliations:** Fischell Department of Bioengineering, University of Maryland, College Park, MD; Department of Neurosurgery, University of Maryland School of Medicine, Baltimore, MD

**Author notes:** Corresponding author: Steven M. Jay, PhD, University of Maryland, 8278 Paint Branch Dr, College Park, MD 20742.

**Keywords:** exosomes, MSCs, GelMA, 3D printing

## Abstract

Extracellular vesicles (EVs) represent an emerging class of therapeutics with significant potential and broad applicability. However, a general limitation is their rapid clearance after administration. Thus, methods to enable sustained EV release are of great potential value. Here, we demonstrate that EVs from mesenchymal stem/stromal cells (MSCs) can be incorporated into 3D-printed gelatin methacrylate (GelMA) hydrogel bioink, and that the initial burst release of EVs can be reduced by increasing the concentration of crosslinker during gelation. Further, the data show that MSC EV bioactivity in an endothelial gap closure assay is retained after the 3D printing and photocrosslinking processes. Our group previously showed that MSC EV bioactivity in this assay correlates with pro-angiogenic bioactivity in vivo, thus these results indicate therapeutic potential of MSC EV-laden GelMA bioinks.

## 1. Introduction

Extracellular vesicles (EVs) derived from mesenchymal stem/stromal cells (MSCs) have been implicated as potential therapeutics in a wide variety of applications [1–12], including in humans [13]. The abilities of MSC EVs to deliver bioactive macromolecular cargo to recipient cells as well as their more drug-like qualities as compared to whole cell-based therapies make MSC EVs especially attractive for tissue repair and regeneration. However, EVs in general are also characterized by relatively rapid clearance from the blood after systemic delivery [14–16], limiting the duration of their therapeutic effects. Thus, approaches to enable sustained delivery of MSC EVs have been investigated to enhance their efficacy.

In particular, hydrogel-based systems for sustained EV release have shown utility. EVs have been integrated into hydrogels by electrostatic or biochemically-mediated attachment or via simple mixing, with ionic crosslinking and photopolymerization used to further restrict EV diffusion and prolong release [17]. For example, hydrogel patches created via gelation of collagen within a gelfoam mesh enabled sustained release of induced pluripotent stem cell-derived cardiomyocyte EVs, resulting in improved recovery from myocardial infarction in rats [18]. Further, injectable self-healing hydrogels have also been shown to allow for sustained MSC EV release to promote bone repair [19] and recovery from spinal cord injury [20]. Hydrogels are also useful as bioinks for 3D-printing, enabling reproducible and custom-designed therapeutic interventions [21]. EVs have been shown to be compatible with 3D-bioprinting [22], and MSC EVs specifically have been integrated into bioprinted hydrogels to promote osteochondral repair [23] as well as neurogenesis in a spinal cord injury model [24]. However, reports of release profiles and bioactivity of MSC EVs from bioprinted constructs remain scarce.

Here, we report the incorporation of MSC EVs into a gelatin methacrylate (GelMA) bioink for the purposes of sustained release. We demonstrate that the initial release rate can be controlled via varying photoinitiator concentration and that MSC EV bioactivity in an endothelial gap closure assay is retained after bioprinting and photopolymerization. These results inform the use of EV-laden bioinks for therapeutic applications.

## 2. Materials and Methods

### 2.1 Cell culture

Bone marrow-derived mesenchymal stem cells (BDMSCs) were purchased from ATCC. Cells initially seeded were designated as passage 1 (P1). BDMSCs were plated in a T175 polystyrene flask and cultured in BDMSC media, consisting of Dulbecco’s Modified Eagle’s Medium (DMEM), supplemented with 10% fetal bovine serum (FBS), 1% non-essential amino acids, and 1% penicillin/streptomycin. Cells were grown to 80-90% confluency and replated at 250,000 cells/T175 flask until P4 when the cells were stored in liquid nitrogen within media supplemented with 10% DMSO as a cryoprotectant for all experimental purposes. Human umbilical vein endothelial cells (HUVECs) were purchased from Promocell and cell passage was again designated as P1. HUVECs were cultured in Endothelial Growth Medium-2 (EGM) (Promocell, C-22221 and C-39221). HUVECs and BDMSCs were repeatedly grown to 80-90% confluency and replated at 500,000 cells/T75 flask and 250,000 cells/T175 flask, repeatedly, until P4 when the cells were stored in liquid nitrogen in media supplemented with 10% of DMSO as a cryoprotectant for all experimental purposes. For EV collection, BDMSCs were cultured in EV-depleted FBS media. FBS (VWR) was heat inactivated (HI) at 56°C for 30 min with intermittent bottle inversions, and then chilled on ice for 30 min. HI-FBS was then centrifuged at 100,000 x *g* for at least 16 h and the supernatant was filtered through a 0.20 μm bottle top filter (VWR) for subsequent use in BDMSC media.

### 2.2 EV isolation and characterization

After day 6 of collecting EV-rich media (120 mL), EVs were isolated using multiple rounds of centrifugation. First, media was centrifuged at 1000 x *g* for 10 min to remove any cells that had detached from the flask. The supernatant from this centrifugation was then centrifuged at 2000 x *g* to remove any large debris. The following supernatant was then centrifuged at 10,000 x *g* for 30 min to remove smaller debris. Finally, to collect the EVs, the supernatant was centrifuged at 100,000 x *g* for 2 h in 8 tubes, each capable of holding approximately 26 mL. The supernatant was discarded, and the remaining EV pellets were resuspended in sterile PBS (500 μL), transferring the PBS from each of the 8 tubes. EVs were washed 3X with sterile PBS using four Nanosep Centrifugal Devices with 300 kDa MWC Omega Membranes (Pall, OD300C35). EVs were resuspended in PBS (150 μL) and stored at −20°C for up to one month, with less than 2 freeze/thaw cycles.

Negative staining of EVs was performed to obtain transmission electron microscopy (TEM) images. Briefly, 4% paraformaldehyde (20 μL) was added to washed EVs (15 μL). The paraformaldehyde/EV solution was placed on parafilm and a resulting droplet formed. A carbon film grid was placed on the droplet for 20 min. The grid was gently picked up with tweezers and excess liquid was blotted off by touching the side at a 45°C Whatman paper. The grid was then transferred to sterile PBS to wash before floating on a droplet (50 μL) of 1% glutaraldehyde for 5 min.

Molecular characterization of EVs was accomplished by immunoblot. Approximately 100 µg of EVs and 25 µg of BDMSC lysates determined by protein content (BCA assay) were used for immunoblots. EV markers CD63, TSG101, and Alix as well as endoplasmic reticulum marker Calnexin were assessed according to standards laid out by the International Society for Extracellular Vesicles [25]. Primary antibodies for CD63 (Proteintech, 25682-1-AP), TSG101 (Abcam, ab125011), Calnexin (Cell Signaling, C5C9), and ALIX (Abcam, ab186429) were used at 1:1000 dilution. Secondary goat anti-rabbit IRDye 800CW (LICOR, 925–32210) was used at a dilution of 1:10,000. Bands were detected with a LI-COR Odyssey CLX Imager.

### 2.3 EV bioactivity assessment

To assess EV activity, an *in vitro* endothelial gap closure assay was used. P4 HUVECs were seeded in a gelatin-coated 24 well plate (500 μL total volume/well) at 200,000 cells/well in EGM2 and allowed to grow until a uniform monolayer was formed (24 h). Medium was replaced with endothelial basal media (EBM) (Promocell, C-22221) supplemented with 0.5% EV-depleted FBS for 24 h. The cell monolayer was then scratched using a 200 μL pipette tip (Rainin, 30389243). Media was then replaced using the same serum starving conditions, but this time supplemented with 100 μg/mL of respective EVs based on BCA quantification of EV-associate protein. EV-depleted EGM and EV-depleted EBM were used as positive and negative controls, respectively. In studies with 0.1% LAP and 0.2% LAP, 100 μg/mL of pooled GelMA was used as a GelMA control. In the study with 0.2% LAP, pooled EVs released from GelMA between hour 1 and hour 24 as well as pooled EVs released between hour 24 and day 7 were also investigated by treating the scratch with 100 μg/mL EVs based on BCA. As controls, pooled EVs suspended at room temperature incubating at 37°C between hour 1 and hour 24 as well as between hour 24 and day 7 were used to treat the cell gap with 100 μg/mL. Lastly, stock EVs stored at −20°C throughout the duration of the release experiment were also used as a positive control at 100 μg/mL. The cell gap was imaged at 0 h and 12 h. Overall gap closure was determined as the percentage of area covered by endothelial cells versus the gap remaining after the latter time point image using ImageJ as previously described [16].

### 2.4 GelMA synthesis, hydrogel preparation and characterization

Gelatin methacrylate was synthesized from type A gelatin (Sigma Aldrich, G2500). Type A gelatin was first dissolved in PBS at 50°C for 20 min at 10% w/v. Then, 0.8 g of methacrylic anhydride (MA) (Sigma Aldrich, 276685) per every 1 g of gelatin in solution was added dropwise to the gelatin solution with rigorous stirring, while still maintaining a reaction temperature of 50°C. The reaction was left to continue for another 1.5 h at 50°C. After this, the contents were transferred to a 50 mL tube and centrifuged at 1000 x *g* for 2 min to remove excess MA. The supernatant was diluted 1:1 with PBS and dialyzed against deionized water across a 10 kDa molecular weight cutoff (ThermoFisher, 66830) for at least 3 days at a temperature of 50°C. The final product was then transferred to a 50 mL tube, frozen overnight at −80°C, and then lyophilized for at least 5 d. The final product was stored in a 50 mL tube with parafilm over its cap at room temperature.

GelMA hydrogels were created from 7% w/v of GelMA in PBS. Photoinitiator lithium phenyl-2,4,6-trimethylbenzoylphosphinate (LAP) (Tocris, 6146) at concentrations of 0.1%, 0.2%, and 0.4% w/v of LAP to GelMA solution were tested. Once LAP was added, the solution was exposed to ultraviolet light for 1 min to initiate the crosslinking process and form the hydrogel. For initial GelMA characterization studies, a circular disc mold was used to create 1mL GelMA hydrogels. Hydrogels were frozen at −80°C and then lyophilized overnight for scanning electron microscopy (SEM). Additional GelMA hydrogels were created for rheological testing (ARES-G2, TA Instruments). Hydrogels were inserted between to flat parallel plates and underwent a range of oscillation frequencies between 0.1 to 10 rad/s with a constant strain of 0.1%. Storage module and loss modulus were plotted over this frequency sweep.

### 2.5 3D-printing GelMA constructs

All 3D-printed GelMA constructs were printed with 7% w/v GelMA in PBS. EV-loaded GelMA discs were printed with doped GelMA bioink with an EV concentration (based on BCA) of 8.84 µg EVs/ µL. The appropriate amount LAP was added to each bioink for each respective study. The final solution of each bioink was vortexed and incubated at 37°C until all GelMA was fully dissolved. The solution was then transferred to a 30 mL tin foil-wrapped amber printer barrel. The piston was added and then the solution was inverted and uncapped to release any visible bubbles. The barrel with bioink was then centrifuged at 1000 x *g* for 5 min to eliminate smaller air bubbles. The bioink underwent brief flash freezing on ice for 5 min following centrifugation and then placed in the ambient UV tool on a BioAssemblyBot (Advanced Solutions, Louisville, KY). A 22-gauge needle was then added to the barrel and a petri dish was added to the printer stage as the collection apparatus. Homogenous discs were printed with a diameter of 6 mm and a thickness of 2 mm. GelMA control discs were printed with the GelMA solution alone and designed using the TSIM Software associated with the BioAssemblyBot. For 7% GelMA, the settings were as followed: pressure: 30 psi; speed: 2 mm/s; UV Curing: print a layer, cure a layer; cure speed: 20 mm/s; light irradiance: 850 mW/cm^2^. Gels were partially crosslinked during printing, and then fully crosslinked with 1 min of UV crosslinking in a UV box. Final gels were covered with PBS for 1 min to aid in lifting the same off the petri dish. PBS was discarded and gels were transferred to a new 48-well plate for release studies.

### 2.6 EV release studies

All EV release studies were performed in 48-well plates in triplicate. After the initial time spent in PBS for 1 min to remove the GelMA discs from the printing surface, discs were placed in their respective wells of a 48 well dish and 1.7 mL of PBS was placed in each well at 37°C. For release of the EV-loaded GelMA with 0.1% LAP, PBS was collected and replaced with 1.7 mL of fresh PBS on hours 1, 4, and 8 as well as days 1, 2, 3, 4, 7, 14, and 21. For release studies of EV-loaded GelMA with 0.2% LAP, PBS was collected and replaced with 1.7 mL of fresh PBS on hours 1, 4, and 8 as well as days 1, 3, 7, 14, and 21. EVs within each of the 1.7 mL PBS collected from the time points were concentrated using 300 kDa MWC Omega Membranes (Pall, OD300C35) in which PBS was removed via centrifugation at 8000 x *g* for 20 min. Then, all samples were resuspended in 60 µL of PBS for quantification. Quantification of EV release was accomplished using an ExoELISA-ULTRA Complete Kit (CD63 Detection; System Biosciences, Mountain View, CA). Briefly, washed EVs were combined with the kit’s Coating Buffer and added to a 96-well plate. The plate was left to incubate for 1 h at 37°C in order for the EVs to bind to the surface. The plate was then washed three times for 5 min each using the wash buffer the kit provided. After this, the plate was incubated with the primary CD63 antibody (1:100 dilution) at room temperature for 1 h under gentle shaking. Plates were again washed 3X for 5 min each with the Wash Buffer. The secondary antibody provided was added (1:5000 dilution) at room temperature for 1 h. The plates were washed and incubated with the kit’s Supersensitive TMP ELISA Substrate at room temperature for 15 min, and the reaction was terminated using the kit’s Stop Buffer Solution. Absorbance was measured at 450 nm using a plate reader. The number of CD63+ EVs/mL was obtained using an exosomal CD63 standard curve produced with the kit’s exosome standards. The total number of EVs was calculated based on the total volume of EVs initially collected.

### 2.7 Statistical analyses

All analyses were performed in triplicate. Data are presented as mean□±□SEM. One-way ANOVAs with Holm-Sidak’s multiple comparison tests were used to determine statistical significance in *in vitro* scratch assays. Student’s t-tests were used to determine statistical significance of EV release on individual days. All statistical analysis was performed with Prism 7 (GraphPad Software, La Jolla, CA). Notation for significance in figures are as follows: ns = P□>□0.05, * = P□<□0.05; ** = P□<□0.01; *** = P□<□0.001; **** = P□<□0.0001.

## 3. Results

### 3.1 MSC EV characterization and in vitro bioactivity

Particle size analyses for MSC EV samples were conducted using nanotracking analysis (NTA), revealing a hydrodynamic diameter range between 30 nm and 250 nm (**Figure 1A**). Immunoblot analyses confirmed the presence of EV-associated proteins CD63, TSG101, Alix, and Flotillin-1 as well as the absence of endoplasmic reticulum marker Calnexin in the EV isolate (**Figure 1B)**. EVs were initially screened for bioactivity using an endothelial gap closure assay, as MSC EV characteristics vary by donor [9], [10]. The gap closure assay was conducted using human umbilical vein endothelial cells (HUVECs) that were treated with growth media only (EGM; positive control), basal media only (EBM), or 100 μg/mL MSC EVs in basal media. Brightfield images were taken at time 0 h and 12 h after gap formation. Representative images for each group and timepoint are displayed, with the gap area outlined (**Figure 1C**). Statistical significance was calculated with one-way ANOVA with Holm-Sidak’s test (***p<0.001, **p<0.01) (n=3) (**Figure 1D**).

**Figure 1:**
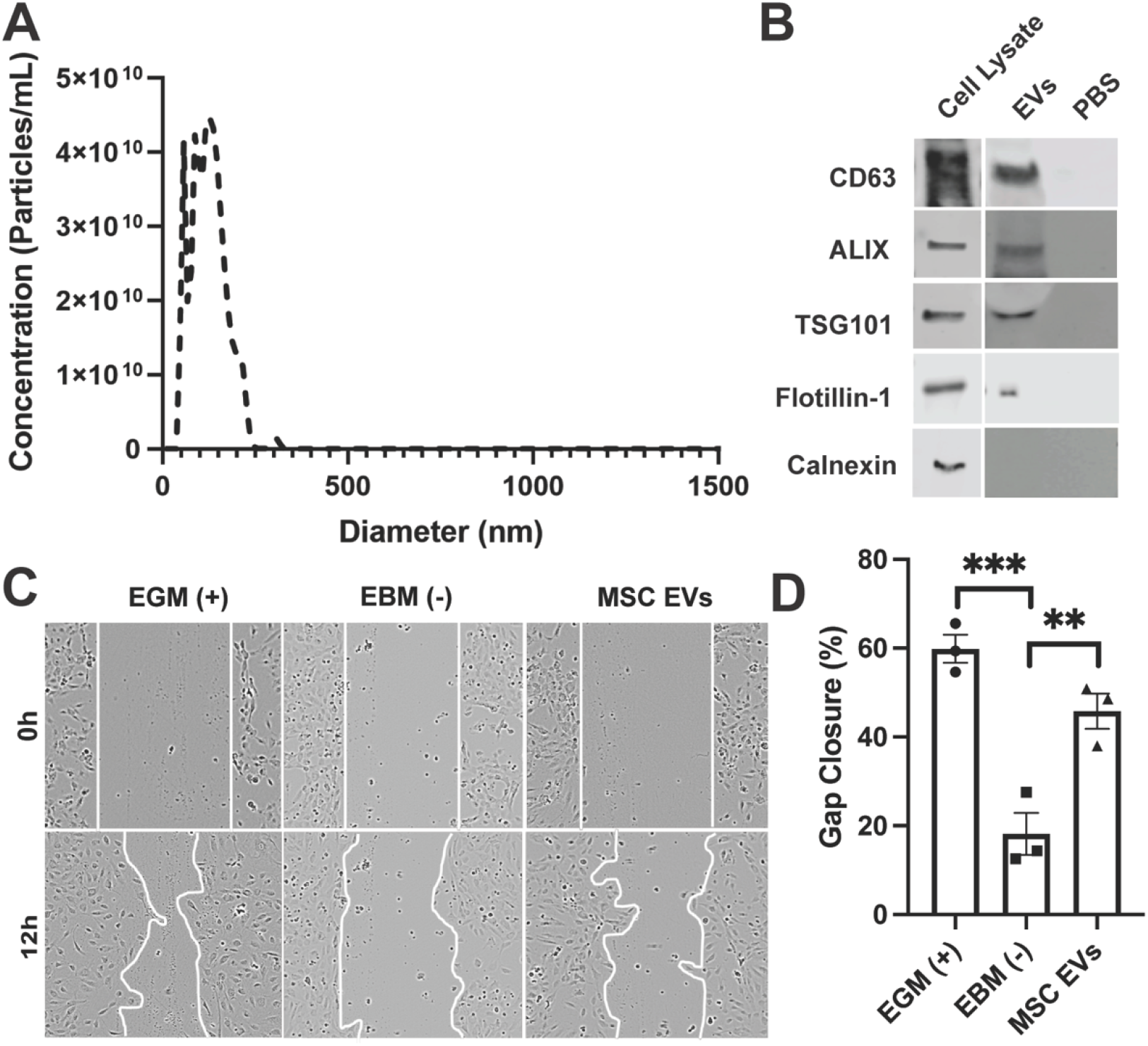
EV characterization and bioactivity assessment. (A) Nanotracking analysis of MSC EVs. (B) Immunoblots of EV-associated markers CD63, ALIX, TSG101, and Flotillin-1 as well as endoplasmic reticulum (ER) marker (Calnexin). (C, D) HUVECs were treated with growth media only (EGM; positive control), basal media only (EBM; negative control), 100 μg/mL MSC EVs in basal media. Images were taken at time 0 (brightfield) and 12h post wounding of the cell layer. Representative images for each group and timepoint are displayed and the gap area outlined with a white line. Statistical significance was calculated using one-way ANOVA with Holm Sidak’s test (**p<0.01, ***p<0.001) (n=3).

### 3.2 Synthesis and characterization of GelMA Hydrogels

Gelatin methacrylate (GelMA) was synthesized from type A gelatin reacted with methacrylic anhydride at 50°C (**Figure 2A**). Verification of GelMA synthesis was confirmed by proton NMR (**Figure 2B**). A proton NMR of gelatin is shown in **Figure 2B** on the left. A proton NMR of GelMA can be seen in **Figure 2B** on the right in which a doublet of doublets peak can be visualized (box) corresponding to the vinyl group from the methacrylate functionalized to the amino terminal of lysine within the gelatin. To construct 3D GelMA hydrogels, 7% (w/v) gelatin with either 0.1%, 0.2%, or 0.4% (w/v) lithium phenyl-2,4,6-trimethylbenzoylphosphinate (LAP) were added to circular molds. The molds containing the GelMA/LAP mixture were then exposed to ultraviolet (UV) light for 1 minute to chemically crosslink the GelMA chains, as described previously [26]. The crosslinked discs were then lyophilized (**Figure 2C)** before imaging via scanning electron microscopy, which revealed generally larger pores for GelMA with 0.1% LAP and smaller pore sizes for 0.2% and 0.4% LAP (**Figure 2D**). Mechanical properties were determined using rheometry (**Figure 2E**), with hydrogels crosslinked with 0.2% and 0.4% LAP having storage moduli across the entire frequency range compared to those crosslinked with 0.1% LAP **(Figure 2F**). There was not a considerable difference in loss modulus properties for any of the conditions tested (**Figure 2G**).

**Figure 2:**
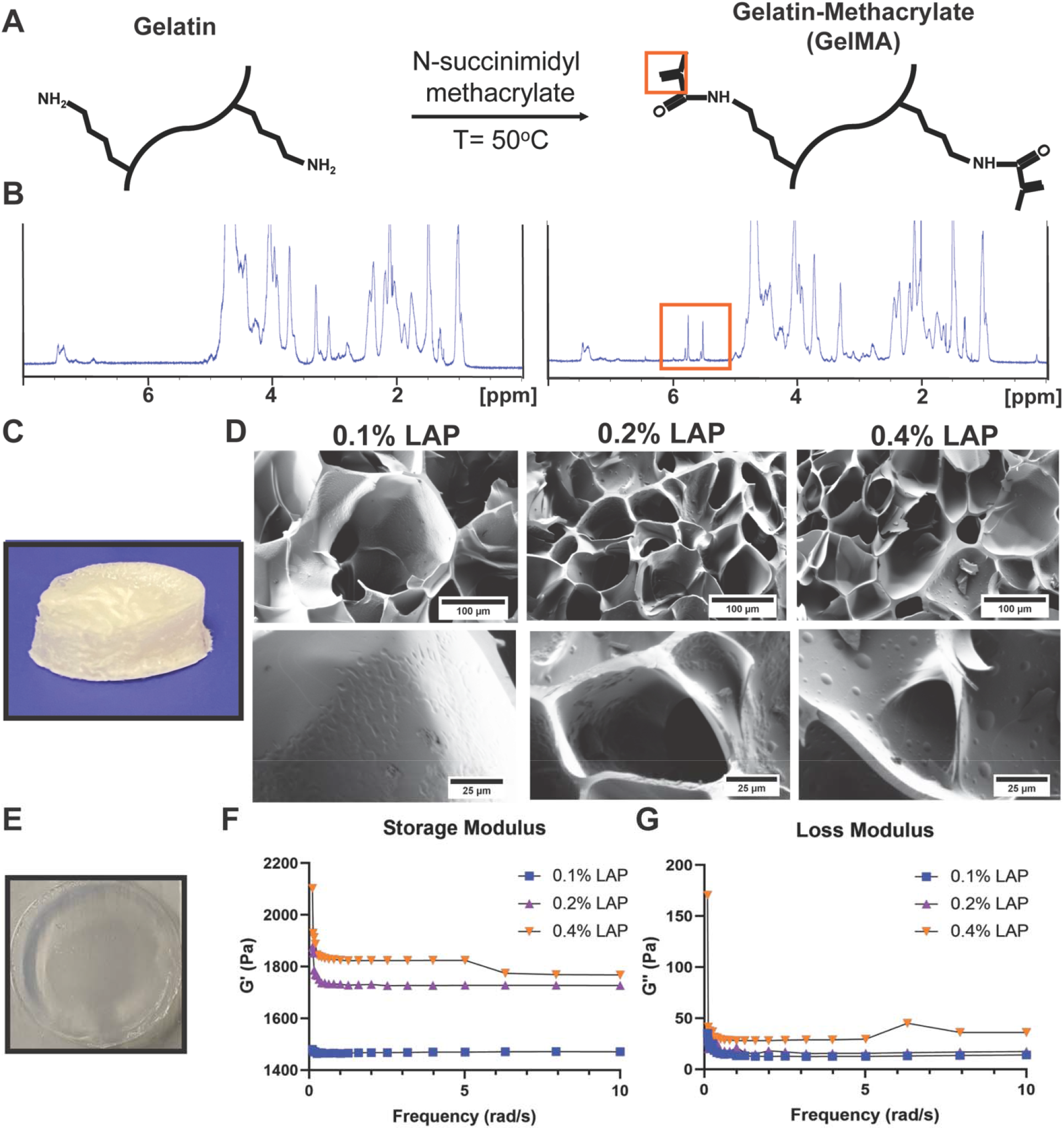
GelMA synthesis and hydrogel characterization. (A) Reaction used to convert gelatin to gelatin methacrylate (GelMA); orange box highlighting vinyl group used in verification of GelMA synthesis *via* NMR. (B) (Left column) Proton NMR of type A gelatin; (right column) proton NMR of GelMA with representative doublet of doublet peak from vinyl group in GelMA(orange box). (C) Lyophilized GelMA disc used for scanning electron microscopy (SEM) imaging. (D) SEM images of 7% (w/v) GelMA with LAP concentrations of 0.1%, 0.2%, and 0.4% (w/v). (E) Swollen hydrogel used for rheometry studies. (F) Storage modulus and (G) loss modulus were measured over a 0-10 rad/s frequency sweep by rheometry (n=3).

### 3.3 MSC EV incorporation into and release from GelMA bioink

To assess whether MSC EV bioactivity is affected by the 3D-printing process, GelMA bioink was made by suspending 8.84 μg EVs/μL in 7% GelMA in phosphate buffered saline (PBS) with either 0.1%, 0.2%, or 0.4% LAP, resulting in 500 μg of EVs per GelMA disc with a 6 mm diameter and 2 mm thickness (**Figure 3A**). The 0.4% LAP constructs exhibited premature photocrosslinking that prevented successful printing, and thus this condition was not pursued further. Successfully printed constructs were placed in 1.7 mL of PBS and samples were collected at indicated time points to assess EV release by CD63 ELISA. Gel crosslinked with 0.1% LAP displayed a significant burst release, while the release from 0.2% LAP crosslinked gels was more prolonged over the first 3 days (**Figure 3B**). In both cases, release was essentially complete by 14 d. Bioactivity of EVs collected separately at 1 d post-incubation was assessed by endothelial gap closure assay and found to be significant for both gel constructs assessed (**Figure 4**), indicating that MSC EVs can retain activity for sustained release applications.

**Figure 3:**
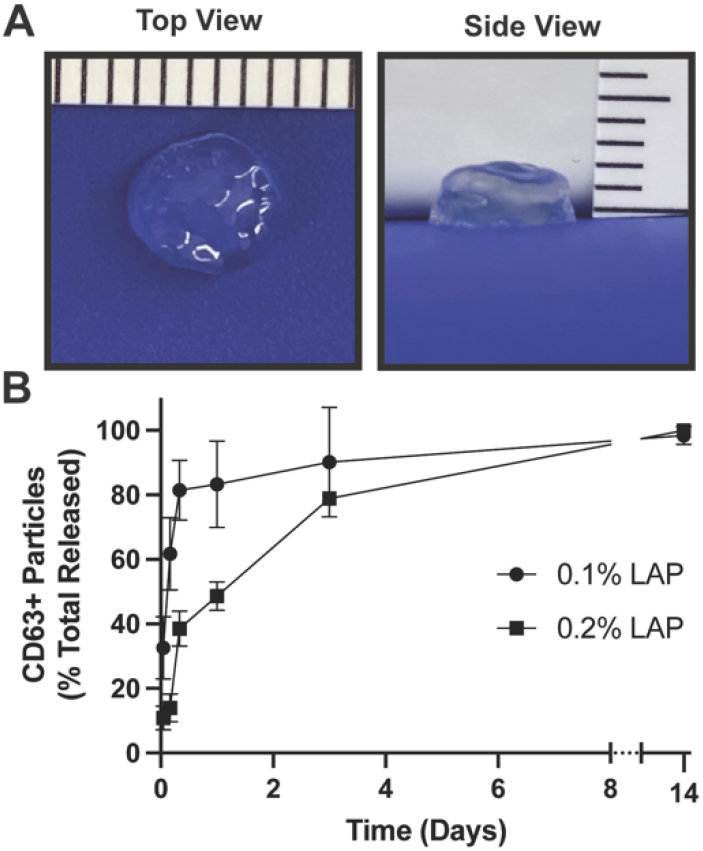
EV release from 3D-printed GelMA hydrogels is affected by crosslinking. (A) 3D-printed GelMA constructs showing accurate 6 mm diameter and 2 mm thickness intended for printing. (B) Release profile of EVs over time determined by CD63+ particles identification by ELISA.

**Figure 4:**
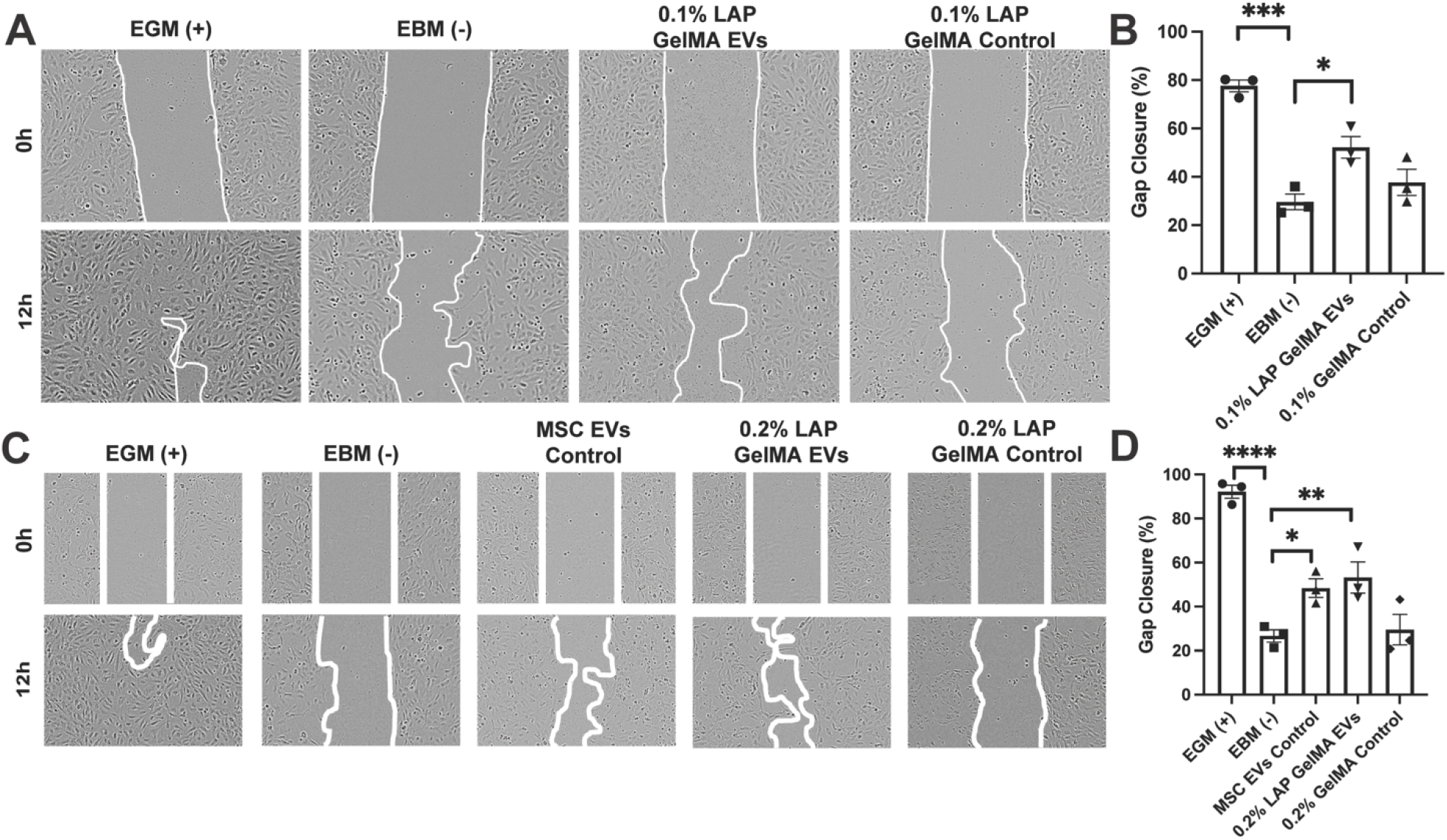
Bioactivity of MSC EVs released from 3D-printed GelMA hydrogels. HUVECs were treated with growth media only (EGM; positive control), basal media only (EBM; negative control), and 100 μg/mL of pooled EVs collected between hour 1 and hour 24 from the indicated constructs. GelMA only controls were included. (A, B) Results of EVs released from GelMA hydrogels crosslinked with 0.1% LAP. (C,D) Results of EVs released from GelMA hydrogels crosslinked with 0.2% LAP. An additional control of unmodified MSC EVs was included. Representative images for each group and timepoint are displayed with the gap area outlined. Statistical significance was calculated using one-way ANOVA with Holm Sidak’s test (*p<0.05, **p<0.01, ***p<0.001, ****p<0.0001) (n=3).

## 4. Discussion

MSC EVs are widely implicated as therapeutics with anti-inflammatory and pro-angiogenic potential, and thus their incorporation into and sustained release from biocompatible hydrogels has many potential applications. The results of these studies show that MSC EVs can be successfully incorporated into GelMA bioinks, retaining activity in an endothelial gap closure assay that is indicative of angiogenic therapeutic potential [27]. Further, the initial release phase of MSC EVs from GelMA is controllable via crosslinker concentration, providing some versatility depending on the application for which such composite bioinks may be used.

The choice of MSCs as the source cells for EVs in these experiments was made based on their broad therapeutic potential as indicated above, and the MSC ECs utilized here were of the expected composition and activity (**Figure 1**) as indicated in the literature [27, 28]. Hydrogel formulations using a variety of materials have been proposed for EV controlled release in various applications [19, 24, 29, 30]. We chose to investigate GelMA in these studies due to its widespread use as a 3D printable bioink based on its biocompatibility, biodegradability, and amenability to a wide array of chemical modifications. The GelMA synthesized for this work had comparable viscoelasticity to prior reports [31], and the decreased pore size associated with increasing LAP content (**Figure 2**) is also consistent with a previous report showing an inverse relationship between photoinitiator (Irgacure 2959) concentration and GelMA hydrogel porosity [32].

EV release studies were conducted by passively loading MSC EVs in 7% (w/v) GelMA constructs, with initial release able to be modulated based on LAP concentration (**Figure 3**). The release profile of encapsulated cargo in hydrogels is a result of various interactions (i.e. cargo/carrier, cargo/cargo, etc.) and release mechanisms (i.e. diffusion, polymer swelling/degradation, etc.). Previous studies for sustained release of hydrophilic cargo from GelMA formulations suggest diffusion as an important mechanism release *in vitro*. As such, altering hydrogel porosity and mesh architecture by modifying GelMA synthesis/crosslinking parameters significantly affects release profile of encapsulated cargo [33]. Here, increased polymer mesh size and porosity increases effective diffusion rate of loaded cargo, which in turn increases release rate [34]. Overall, the results indicate a key role for diffusion in EV release from GelMA. Burst release is a common phenomenon in hydrogel-based drug carriers, where loosely associated cargo gets liberated immediately after contact with the release medium. After 24 h, we observed little to no detectable CD63+ particles in the 0.1% LAP group. In contrast, a slight majority (∼55%) of loaded CD63+ EVs in the 0.2% LAP group was detected after the initial 24 h, demonstrating the effectiveness of increased crosslinking on limiting EV diffusion. Electrostatic interactions between cargo/GelMA can also affect release profile in some cases. Charged proteins such as basic fibroblast growth factor and bone morphogenic protein-2 maybe sequestered fully GelMA constructs, presumably via adsorption [35]. Other studies have demonstrated that GelMA formulation parameters such as ionic strength and pH can affect cargo loading, GelMA mesh characteristics, and overall net charge [36]. Although EV/GelMA electrostatic interactions were beyond the scope of this study, EVs generally carry a net negative charge and such interactions may be an important factor to consider in future studies.

Critically, the bioactivity of the released MSC EVs was confirmed via endothelial gap closure assay (**Figure 4**). Our results suggest that EVs are likely to remain bioactive through 3D bioprinting and photocrosslinking with LAP, which is critical to their ultimate utility as part of a composite bioink. While further validation is needed, it is expected that EVs from any source cell type could be utilized in a similar fashion, creating therapeutic opportunities for myriad diseases and injuries.

Several limitations of our study should be noted. We chose to utilize 7% w/v of GelMA due to compatibility with 3D printing, however increasing the concentration of GelMA in the bioink could further modify hydrogel porosity as a means to gain further control over the EV release profile [32]. Additional studies testing the limits of viscosity with increased GelMA in solution should be performed. We also used a 3D printer that operates at room temperature, and thus temperature optimization was not attempted. Although diffusion is an important mechanism determining release kinetics, it is currently unknown whether electrostatic EV/GelMA interactions also play a significant role. Moreover, GelMA contains enzymatic degradation sites for collagenases and matrix metalloproteinases (MMPs). Thus, future studies could examine release kinetics in the presence of such enzymes to better model *in vivo* release.

## 5. Conclusion

MSC EVs can be incorporated into 3D-printed, photocrosslinked GelMA constructs without compromising at least one key aspect of their bioactivity. Further, EV burst release can be significantly reduced by increasing the crosslinker (LAP) concentration during gelation. Thus, EV-laden GelMA bioinks merit further exploration for therapeutic applications.

## References

1. Li, Z., F. Liu, X. He, X. Yang, F. Shan, and J. Feng, Exosomes derived from mesenchymal stem cells attenuate inflammation and demyelination of the central nervous system in EAE rats by regulating the polarization of microglia. Int Immunopharmacol, 2019. 67: p. 268–280.

2. Zou, L., X. Ma, S. Lin, B. Wu, Y. Chen, and C. Peng, Bone marrow mesenchymal stem cell-derived exosomes protect against myocardial infarction by promoting autophagy. Exp Ther Med, 2019. 18(4): p. 2574–2582.

3. Sun, X., A. Shan, Z. Wei, and B. Xu, Intravenous mesenchymal stem cell-derived exosomes ameliorate myocardial inflammation in the dilated cardiomyopathy. Biochem Biophys Res Commun, 2018. 503(4): p. 2611–2618.

4. Fujii, S., Y. Miura, A. Fujishiro, T. Shindo, Y. Shimazu, H. Hirai, H. Tahara, A. Takaori-Kondo, T. Ichinohe, and T. Maekawa, Graft-Versus-Host Disease Amelioration by Human Bone Marrow Mesenchymal Stromal/Stem Cell-Derived Extracellular Vesicles Is Associated with Peripheral Preservation of Naive T Cell Populations. Stem Cells, 2018. 36(3): p. 434–445.

5. Long, Q., D. Upadhya, B. Hattiangady, D.K. Kim, S.Y. An, B. Shuai, D.J. Prockop, and A.K. Shetty, Intranasal MSC-derived A1-exosomes ease inflammation, and prevent abnormal neurogenesis and memory dysfunction after status epilepticus. Proc Natl Acad Sci U S A, 2017. 114(17): p. E3536–E3545.

6. Ophelders, D.R., T.G. Wolfs, R.K. Jellema, A. Zwanenburg, P. Andriessen, T. Delhaas, A.K. Ludwig, S. Radtke, V. Peters, L. Janssen, B. Giebel, and B.W. Kramer, Mesenchymal Stromal Cell-Derived Extracellular Vesicles Protect the Fetal Brain After Hypoxia-Ischemia. Stem Cells Transl Med, 2016. 5(6): p. 754–63.

7. Wang, B., K. Yao, B.M. Huuskes, H.H. Shen, J. Zhuang, C. Godson, E.P. Brennan, J.L. Wilkinson-Berka, A.F. Wise, and S.D. Ricardo, Mesenchymal Stem Cells Deliver Exogenous MicroRNA-let7c via Exosomes to Attenuate Renal Fibrosis. Mol Ther, 2016. 24(7): p. 1290–301.

8. Wang, X., H. Gu, D. Qin, L. Yang, W. Huang, K. Essandoh, Y. Wang, C.C. Caldwell, T. Peng, B. Zingarelli, and G.C. Fan, Exosomal miR-223 Contributes to Mesenchymal Stem Cell-Elicited Cardioprotection in Polymicrobial Sepsis. Sci Rep, 2015. 5: p. 13721.

9. Doeppner, T.R., J. Herz, A. Gorgens, J. Schlechter, A.K. Ludwig, S. Radtke, K. de Miroschedji, P.A. Horn, B. Giebel, and D.M. Hermann, Extracellular Vesicles Improve Post-Stroke Neuroregeneration and Prevent Postischemic Immunosuppression. Stem Cells Transl Med, 2015. 4(10): p. 1131–43.

10. Lai, R.C., F. Arslan, M.M. Lee, N.S. Sze, A. Choo, T.S. Chen, M. Salto-Tellez, L. Timmers, C.N. Lee, R.M. El Oakley, G. Pasterkamp, D.P. de Kleijn, and S.K. Lim, Exosome secreted by MSC reduces myocardial ischemia/reperfusion injury. Stem Cell Res, 2010. 4(3): p. 214–22.

11. Li, Q., Y. Xu, K. Lv, Y. Wang, Z. Zhong, C. Xiao, K. Zhu, C. Ni, K. Wang, M. Kong, X. Li, Y. Fan, F. Zhang, Q. Chen, Y. Li, Q. Li, C. Liu, J. Zhu, S. Zhong, J. Wang, Y. Chen, J. Zhao, D. Zhu, R. Wu, J. Chen, W. Zhu, H. Yu, R. Ardehali, J.J. Zhang, J. Wang, and X. Hu, Small extracellular vesicles containing miR-486-5p promote angiogenesis after myocardial infarction in mice and nonhuman primates. Sci Transl Med, 2021. 13(584).

12. Park, K.S., K. Svennerholm, G.V. Shelke, E. Bandeira, C. Lasser, S.C. Jang, R. Chandode, I. Gribonika, and J. Lotvall, Mesenchymal stromal cell-derived nanovesicles ameliorate bacterial outer membrane vesicle-induced sepsis via IL-10. Stem Cell Res Ther, 2019. 10(1): p. 231.

13. Kordelas, L., V. Rebmann, A.K. Ludwig, S. Radtke, J. Ruesing, T.R. Doeppner, M. Epple, P.A. Horn, D.W. Beelen, and B. Giebel, MSC-derived exosomes: a novel tool to treat therapy-refractory graft-versus-host disease. Leukemia, 2014. 28(4): p. 970–3.

14. Matsumoto, A., Y. Takahashi, H.Y. Chang, Y.W. Wu, A. Yamamoto, Y. Ishihama, and Y. Takakura, Blood concentrations of small extracellular vesicles are determined by a balance between abundant secretion and rapid clearance. J Extracell Vesicles, 2020. 9(1): p. 1696517.

15. Mirzaaghasi, A., Y. Han, S.H. Ahn, C. Choi, and J.H. Park, Biodistribution and Pharmacokinectics of Liposomes and Exosomes in a Mouse Model of Sepsis. Pharmaceutics, 2021. 13(3).

16. Yamamoto, A., Y. Yasue, Y. Takahashi, and Y. Takakura, Determining the role of surface glycans in the pharmacokinetics of small extracellular vesicles. J Pharm Sci, 2021.

17. Huang, J., J. Xiong, L. Yang, J. Zhang, S. Sun, and Y. Liang, Cell-free exosome-laden scaffolds for tissue repair. Nanoscale, 2021. 13(19): p. 8740–8750.

18. Liu, B., B.W. Lee, K. Nakanishi, A. Villasante, R. Williamson, J. Metz, J. Kim, M. Kanai, L. Bi, K. Brown, G. Di Paolo, S. Homma, P.A. Sims, V.K. Topkara, and G. Vunjak-Novakovic, Cardiac recovery via extended cell-free delivery of extracellular vesicles secreted by cardiomyocytes derived from induced pluripotent stem cells. Nat Biomed Eng, 2018. 2(5): p. 293–303.

19. Wang, L., J. Wang, X. Zhou, J. Sun, B. Zhu, C. Duan, P. Chen, X. Guo, T. Zhang, and H. Guo, A New Self-Healing Hydrogel Containing hucMSC-Derived Exosomes Promotes Bone Regeneration. Front Bioeng Biotechnol, 2020. 8: p. 564731.

20. Wang, C., M. Wang, K. Xia, J. Wang, F. Cheng, K. Shi, L. Ying, C. Yu, H. Xu, S. Xiao, C. Liang, F. Li, B. Lei, and Q. Chen, A bioactive injectable self-healing anti-inflammatory hydrogel with ultralong extracellular vesicles release synergistically enhances motor functional recovery of spinal cord injury. Bioact Mater, 2021. 6(8): p. 2523–2534.

21. Santoro, M., J. Navarro, and J.P. Fisher, Micro-and Macrobioprinting: Current Trends in Tissue Modeling and Organ Fabrication. Small Methods, 2018. 2(3).

22. Yerneni, S.S., T.L. Whiteside, L.E. Weiss, and P.G. Campbell, Bioprinting exosome-like extracellular vesicle microenvironments. Bioprinting, 2019. 13: p. e00041.

23. Chen, P., L. Zheng, Y. Wang, M. Tao, Z. Xie, C. Xia, C. Gu, J. Chen, P. Qiu, S. Mei, L. Ning, Y. Shi, C. Fang, S. Fan, and X. Lin, Desktop-stereolithography 3D printing of a radially oriented extracellular matrix/mesenchymal stem cell exosome bioink for osteochondral defect regeneration. Theranostics, 2019. 9(9): p. 2439–2459.

24. Cheng, J., Z. Chen, C. Liu, M. Zhong, S. Wang, Y. Sun, H. Wen, and T. Shu, Bone mesenchymal stem cell-derived exosome-loaded injectable hydrogel for minimally invasive treatment of spinal cord injury. Nanomedicine (Lond), 2021. 16(18): p. 1567–1579.

25. Thery, C., K.W. Witwer, E. Aikawa, M.J. Alcaraz, J.D. Anderson, R. Andriantsitohaina, A. Antoniou, T. Arab, F. Archer, G.K. Atkin-Smith, D.C. Ayre, J.M. Bach, D. Bachurski, H. Baharvand, L. Balaj, S. Baldacchino, N.N. Bauer, A.A. Baxter, M. Bebawy, C. Beckham, Bedina Zavec, A. Benmoussa, A.C. Berardi, P. Bergese, E. Bielska, C. Blenkiron, S. Bobis-Wozowicz, E. Boilard, W. Boireau, A. Bongiovanni, F.E. Borras, S. Bosch, C.M. Boulanger, X. Breakefield, A.M. Breglio, M.A. Brennan, D.R. Brigstock, A. Brisson, M.L. Broekman, J.F. Bromberg, P. Bryl-Gorecka, S. Buch, A.H. Buck, D. Burger, S. Busatto, D. Buschmann, B. Bussolati, E.I. Buzas, J.B. Byrd, G. Camussi, D.R. Carter, S. Caruso, L.W. Chamley, Y.T. Chang, C. Chen, S. Chen, L. Cheng, A.R. Chin, A. Clayton, S.P. Clerici, A. Cocks, E. Cocucci, R.J. Coffey, A. Cordeiro-da-Silva, Y. Couch, F.A. Coumans, B. Coyle, R. Crescitelli, M.F. Criado, C. D’Souza-Schorey, S. Das, A. Datta Chaudhuri, P. de Candia, E.F. De Santana, O. De Wever, H.A. Del Portillo, T. Demaret, S. Deville, A. Devitt, B. Dhondt, D. Di Vizio, L.C. Dieterich, V. Dolo, A.P. Dominguez Rubio, M. Dominici, M.R. Dourado, T.A. Driedonks, F.V. Duarte, H.M. Duncan, R.M. Eichenberger, K. Ekstrom, S. El Andaloussi, C. Elie-Caille, U. Erdbrugger, J.M. Falcon-Perez, F. Fatima, J.E. Fish, M. Flores-Bellver, A. Forsonits, A. Frelet-Barrand, F. Fricke, G. Fuhrmann, S. Gabrielsson, A. Gamez-Valero, C. Gardiner, K. Gartner, R. Gaudin, Y.S. Gho, B. Giebel, C. Gilbert, M. Gimona, I. Giusti, D.C. Goberdhan, A. Gorgens, S.M. Gorski, D.W. Greening, J.C. Gross, A. Gualerzi, G.N. Gupta, D. Gustafson, A. Handberg, R.A. Haraszti, P. Harrison, H. Hegyesi, A. Hendrix, A.F. Hill, F.H. Hochberg, K.F. Hoffmann, B. Holder, H. Holthofer, B. Hosseinkhani, G. Hu, Y. Huang, V. Huber, S. Hunt, A.G. Ibrahim, T. Ikezu, J.M. Inal, M. Isin, A. Ivanova, H.K. Jackson, S. Jacobsen, S.M. Jay, M. Jayachandran, G. Jenster, L. Jiang, S.M. Johnson, J.C. Jones, A. Jong, T. Jovanovic-Talisman, S. Jung, R. Kalluri, S.I. Kano, S. Kaur, Y. Kawamura, E.T. Keller, D. Khamari, E. Khomyakova, A. Khvorova, P. Kierulf, K.P. Kim, T. Kislinger, M. Klingeborn, D.J. Klinke, 2nd, M. Kornek, M.M. Kosanovic, A.F. Kovacs, E.M. Kramer-Albers, S. Krasemann, M. Krause, I.V. Kurochkin, G.D. Kusuma, S. Kuypers, S. Laitinen, S.M. Langevin, L.R. Languino, J. Lannigan, C. Lasser, L.C. Laurent, G. Lavieu, E. Lazaro-Ibanez, S. Le Lay, M.S. Lee, Y.X.F. Lee, D.S. Lemos, M. Lenassi, A. Leszczynska, I.T. Li, K. Liao, S.F. Libregts, E. Ligeti, R. Lim, S.K. Lim, A. Line, K. Linnemannstons, A. Llorente, C.A. Lombard, M.J. Lorenowicz, A.M. Lorincz, J. Lotvall, J. Lovett, M.C. Lowry, X. Loyer, Q. Lu, B. Lukomska, T.R. Lunavat, S.L. Maas, H. Malhi, A. Marcilla, J. Mariani, J. Mariscal, E.S. Martens-Uzunova, L. Martin-Jaular, M.C. Martinez, V.R. Martins, M. Mathieu, S. Mathivanan, M. Maugeri, L.K. McGinnis, M.J. McVey, D.G. Meckes, Jr., K.L. Meehan, I. Mertens, V.R. Minciacchi, A. Moller, M. Moller Jorgensen, A. Morales-Kastresana, J. Morhayim, F. Mullier, M. Muraca, L. Musante, V. Mussack, D.C. Muth, K.H. Myburgh, T. Najrana, M. Nawaz, I. Nazarenko, P. Nejsum, C. Neri, T. Neri, R. Nieuwland, L. Nimrichter, J.P. Nolan, E.N. Nolte-’t Hoen, N. Noren Hooten, L. O’Driscoll, T. O’Grady, A. O’Loghlen, T. Ochiya, M. Olivier, A. Ortiz, L.A. Ortiz, X. Osteikoetxea, O. Ostergaard, M. Ostrowski, J. Park, D.M. Pegtel, H. Peinado, F. Perut, M.W. Pfaffl, D.G. Phinney, B.C. Pieters, R.C. Pink, D.S. Pisetsky, E. Pogge von Strandmann, I. Polakovicova, I.K. Poon, B.H. Powell, I. Prada, L. Pulliam, P. Quesenberry, A. Radeghieri, R.L. Raffai, S. Raimondo, J. Rak, M.I. Ramirez, G. Raposo, M.S. Rayyan, N. Regev-Rudzki, F.L. Ricklefs, P.D. Robbins, D.D. Roberts, S.C. Rodrigues, E. Rohde, S. Rome, K.M. Rouschop, A. Rughetti, A.E. Russell, P. Saa, S. Sahoo, E. Salas-Huenuleo, C. Sanchez, J.A. Saugstad, M.J. Saul, R.M. Schiffelers, R. Schneider, T.H. Schoyen, A. Scott, E. Shahaj, S. Sharma, O. Shatnyeva, F. Shekari, G.V. Shelke, A.K. Shetty, K. Shiba, P.R. Siljander, A.M. Silva, A. Skowronek, O.L. Snyder, 2nd, R.P. Soares, B.W. Sodar, C. Soekmadji, J. Sotillo, P.D. Stahl, W. Stoorvogel, S.L. Stott, E.F. Strasser, S. Swift, H. Tahara, M. Tewari, K. Timms, S. Tiwari, R. Tixeira, M. Tkach, W.S. Toh, R. Tomasini, A.C. Torrecilhas, J.P. Tosar, V. Toxavidis, L. Urbanelli, P. Vader, B.W. van Balkom, S.G. van der Grein, J. Van Deun, M.J. van Herwijnen, K. Van Keuren-Jensen, G. van Niel, M.E. van Royen, A.J. van Wijnen, M.H. Vasconcelos, I.J. Vechetti, Jr., T.D. Veit, L.J. Vella, E. Velot, F.J. Verweij, B. Vestad, J.L. Vinas, T. Visnovitz, K.V. Vukman, J. Wahlgren, D.C. Watson, M.H. Wauben, A. Weaver, J.P. Webber, V. Weber, A.M. Wehman, D.J. Weiss, J.A. Welsh, S. Wendt, A.M. Wheelock, Z. Wiener, L. Witte, J. Wolfram, A. Xagorari, P. Xander, J. Xu, X. Yan, M. Yanez-Mo, H. Yin, Y. Yuana, V. Zappulli, J. Zarubova, V. Zekas, J.Y. Zhang, Z. Zhao, L. Zheng, A.R. Zheutlin, A.M. Zickler, P. Zimmermann, A.M. Zivkovic, D. Zocco and E.K. Zuba-Surma, Minimal information for studies of extracellular vesicles 2018 (MISEV2018): a position statement of the International Society for Extracellular Vesicles and update of the MISEV2014 guidelines. J Extracell Vesicles, 2018. 7(1): p. 1535750.

26. Yu, J.R., M. Janssen, B.J. Liang, H.C. Huang, and J.P. Fisher, A liposome/gelatin methacrylate nanocomposite hydrogel system for delivery of stromal cell-derived factor-1alpha and stimulation of cell migration. Acta Biomater, 2020. 108: p. 67–76.

27. Born, L.J., K.H. Chang, P. Shoureshi, F. Lay, S. Bengali, A.T.W. Hsu, S.N. Abadchi, J.W. Harmon, and S.M. Jay, HOTAIR-Loaded Mesenchymal Stem/Stromal Cell Extracellular Vesicles Enhance Angiogenesis and Wound Healing. Adv Healthc Mater, 2021: p. e2002070.

28. Patel, D.B., K.M. Gray, Y. Santharam, T.N. Lamichhane, K.M. Stroka, and S.M. Jay, Impact of cell culture parameters on production and vascularization bioactivity of mesenchymal stem cell-derived extracellular vesicles. Bioeng Transl Med, 2017. 2(2): p. 170–179.

29. Millan, C., L. Prause, Q. Vallmajo-Martin, N. Hensky, and D. Eberli, Extracellular Vesicles from 3D Engineered Microtissues Harbor Disease-Related Cargo Absent in EVs from 2D Cultures. Adv Healthc Mater, 2021: p. e2002067.

30. Han, C., J. Zhou, B. Liu, C. Liang, X. Pan, Y. Zhang, Y. Zhang, Y. Wang, L. Shao, B. Zhu, J. Wang, Q. Yin, X.Y. Yu, and Y. Li, Delivery of miR-675 by stem cell-derived exosomes encapsulated in silk fibroin hydrogel prevents aging-induced vascular dysfunction in mouse hindlimb. Mater Sci Eng C Mater Biol Appl, 2019. 99: p. 322–332.

31. Duchi, S., C. Onofrillo, C.D. O’Connell, R. Blanchard, C. Augustine, A.F. Quigley, R.M.I. Kapsa, P. Pivonka, G. Wallace, C. Di Bella, and P.F.M. Choong, Handheld Co-Axial Bioprinting: Application to in situ surgical cartilage repair. Sci Rep, 2017. 7(1): p. 5837.

32. Benton, J.A., C.A. DeForest, V. Vivekanandan, and K.S. Anseth, Photocrosslinking of gelatin macromers to synthesize porous hydrogels that promote valvular interstitial cell function. Tissue Eng Part A, 2009. 15(11): p. 3221–30.

33. Nguyen, D.K., Y.M. Son, and N.E. Lee, Hydrogel Encapsulation of Cells in Core-Shell Microcapsules for Cell Delivery. Adv Healthc Mater, 2015. 4(10): p. 1537–44.

34. Vigata, M., C. Meinert, S. Pahoff, N. Bock, and D.W. Hutmacher, Gelatin Methacryloyl Hydrogels Control the Localized Delivery of Albumin-Bound Paclitaxel. Polymers (Basel), 2020. 12(2).

35. Samorezov, J.E., E.B. Headley, C.R. Everett, and E. Alsberg, Sustained presentation of BMP-2 enhances osteogenic differentiation of human adipose-derived stem cells in gelatin hydrogels. J Biomed Mater Res A, 2016. 104(6): p. 1387–97.

36. Vigata, M., C. Meinert, N. Bock, B.L. Dargaville, and D.W. Hutmacher, Deciphering the Molecular Mechanism of Water Interaction with Gelatin Methacryloyl Hydrogels: Role of Ionic Strength, pH, Drug Loading and Hydrogel Network Characteristics. Biomedicines, 2021. 9(5).

